# The dCMP deaminase DCTD and the E3 ligase TOPORS are central mediators of decitabine cytotoxicity

**DOI:** 10.1101/2023.12.21.572728

**Authors:** Christopher J. Carnie, Maximilian J. Götz, Chloe S. Palma-Chaundler, Pedro Weickert, Amy R. Wanders, Almudena Serrano-Benitez, Hao-Yi Li, Vipul Gupta, Christian J. Blum, Matylda Sczaniecka-Clift, Guido Zagnoli-Vieira, Giuseppina D’Alessandro, Sean L. Richards, Nadia Gueorguieva, Petra Beli, Julian Stingele, Stephen P. Jackson

## Abstract

The nucleoside decitabine (5-aza-dC) is used to treat several hematological cancers. Upon triphosphorylation and incorporation into DNA, 5-aza-dC induces covalent DNMT1 DNA-protein crosslinks (DPCs) and DNA hypomethylation. However, 5-aza-dC treatment success varies, and relapse is common. Using genome-scale CRISPR/Cas9 screens, we map factors determining 5-aza-dC susceptibility. Unexpectedly, we find that loss of the dCMP deaminase DCTD causes 5-aza-dC resistance, suggesting that 5-aza-dUMP generation underlies most 5-aza-dC cytotoxicity in wild-type cells. Combining results from a subsequent genetic screen in DCTD-deficient cells with identification of the proximal proteome of DNMT1-DPCs, we uncover the ubiquitin/SUMO1 E3 ligase, TOPORS, as a new DPC repair factor. TOPORS is recruited to DNMT1-DPCs in a SUMO-dependent manner and promotes their degradation. Our study suggests that 5-aza-dC-induced DPCs cause cytotoxicity when DPC repair is compromised, while cytotoxicity in wild-type cells arises from perturbed nucleotide metabolism and lays the foundations for the development of predictive biomarkers for decitabine treatment.

## Introduction

Myelodysplastic syndromes (MDS) are a heterogenous group of neoplastic disorders that represent the most common group of hematological malignancies^1^. MDS is characterised by dysplasia and ineffective hematopoiesis, leading to peripheral cytopenia along with a risk of disease progression to acute myelocytic leukemia (AML)^2^. The core therapy for management of MDS consists of the nucleoside analogues 5-azacytidine (5-aza-C) and 5-aza-deoxycytidine (5-aza-dC, also known as decitabine). Decitabine is also used for the treatment of AML and chronic myelocytic leukemia (CML), particularly in elderly patients who are ineligible to undergo more aggressive regimens with agents such as cytarabine (cytosine arabinoside)^3,4^.

5-aza-C and 5-aza-dC are widely believed to exert their therapeutic effects primarily by trapping DNA methyltransferases (DNMTs) onto DNA, thereby inhibiting DNMTs and inducing DNA hypomethylation and re-expression of previously silenced tumor suppressor genes^5,6^. 5-aza-C and 5-aza-dC are thus considered hypomethylating agents (HMAs). Despite their widespread use in treating certain blood cancers, responses to HMAs vary from patient to patient for reasons that have remained unclear^7,8^. Only around 30-50% of patients respond well to HMAs^9,10^, with a subpopulation of patients not responding at all. This is especially problematic because HMAs are given in low doses over long treatment periods of up to six months before the individual treatment effectiveness can be assessed^11^. However, promising new approaches utilising HMAs in combination with other drugs such as the BCL2 inhibitor venetoclax, are emerging^4,12^. Therefore, a detailed understanding of the mechanism(s) of action of HMAs and identification of predictive biomarkers to guide individual therapy is becoming increasingly needed. In addition to DNMT1 DNA-protein crosslink (DPC) induction^13–15^, HMAs also cause broad cytotoxicity by perturbing RNA synthesis and activating immune checkpoints^16^. 5-aza-C, a ribonucleoside, is incorporated into both RNA and DNA, the latter being dependent on reduction of 5-aza-CDP to 5-aza-dCDP by ribonucleotide reductase^17–19^. The deoxyribonucleoside 5-aza-dC is phosphorylated by deoxycytidine kinase (DCK)^20^ to 5-aza-dCMP. 5-aza-dCMP is further phosphorylated by cytidine/uridine monophosphate kinase 1 (CMPK1) and nucleoside diphosphate kinase 1/2 (NME1/2) to generate 5-aza-dCTP, which can be incorporated into nascent DNA^21^. Alternatively, 5-aza-dC or 5-aza-dCMP can be deaminated by dCMP deaminase (DCTD) or cytidine deaminase (CDA) to 5-aza-dU or 5-aza-dUMP, respectively^22,23^. Interestingly, CDA is highly expressed in certain organs, such as the liver and the gut, where it deaminates HMAs. The rapid deamination of HMAs is responsible for their short serum half-life and is believed to result in their detoxification^24,25^. The FDA has recently approved the combination of 5-aza-dC and the CDA inhibitor cedazurine for the treatment of MDS, a strategy aimed at reducing the extent of 5-aza-dC deamination^25^.

Once incorporated into DNA, azacytidines act as a pseudo-substrate for DNMT1 at hemi-methylated CpGs, trapping a covalent DNA-DNMT1 reaction intermediate^5,14^. DPCs are highly toxic DNA lesions that interfere with chromatin-associated processes such as replication and transcription^26,27^. Repair of these lesions requires proteolytic degradation by either the proteasome or dedicated DPC proteases of the Wss1/SPRTN family^28–32^. DPC repair can be initiated in a replication-coupled manner upon polymerase stalling^28–30,33,34^. Additionally, global-genome DPC repair occurs upon DPC SUMOylation by PIAS4 and subsequent ubiquitylation by the SUMO-targeted ubiquitin ligase (STUbL) RNF4, promoting degradation of the DPC by SPRTN and the proteasome^35–38^. How DPC formation or other cytotoxic effects of 5-aza-dC precisely contribute to the overall mode-of-action of HMAs, however, remains unresolved.

Here, we employ a genome-wide CRISPR/Cas9 screen to map genes conferring resistance or sensitivity to decitabine treatment, uncovering a major mode of 5-aza-dC cytotoxicity that acts through its deamination by DCTD. In addition, using a further genome-wide genetic screen with 5-aza-dC in *DCTD* KO cells and a modified iPOND (isolation of proteins on nascent DNA) approach to determine the proximal proteome of DNMT1-DPCs, we categorise the hits from our genetic screens into DPC-dependent and DPC-independent classes. Using this approach, we identify TOPORS, a SUMO1- and ubiquitin E3 ligase, as a decitabine resistance factor that is recruited to decitabine-induced DNMT1-DPCs. TOPORS recruitment is SUMO-dependent but ubiquitin-independent and promotes proteolysis of DNMT1-DPCs. Our findings indicate that 5-aza-dC-induced cytotoxicity is driven by perturbed nucleotide metabolism in wild-type cells and by DNMT1-DPC formation in cells with DPC repair defects.

## Results

### Nucleotide metabolism modulates 5-aza-dC/decitabine cytotoxicity

To profile genetic determinants of 5-aza-dC sensitivity and resistance, we performed a genome-scale CRISPR/Cas9 screen with 5-aza-dC in the CML-derived human cell line HAP1 (Fig 1a). Based on a false discovery rate (FDR) of 0.1, our screen identified 48 genes whose individual loss conferred sensitivity to 5-aza-dC, and 11 genes whose loss conferred resistance (Fig 1b). As expected, inactivation of *SLC29A1* or *DCK* conferred 5-aza-dC resistance, while loss of *SAMHD1* caused sensitization. *SLC29A1* encodes ENT1, a nucleoside transporter that mediates cellular uptake of 5-aza-dC (Fig 1c)^4,39–42^. DCK phosphorylates 5-aza-dC and is thus required for its subsequent incorporation into DNA (Fig. 1c). SAMHD1 is a hydrolase that cleaves dNTPs into deoxynucleosides and triphosphate and has previously been shown to hydrolyse 5-aza-dCTP^43^ (Fig 1c). Therefore, loss of SAMHD1 is expected to increase incorporation of 5-aza-dCTP, thereby increasing cellular sensitivity to 5-aza-dC.

**Figure 1.**
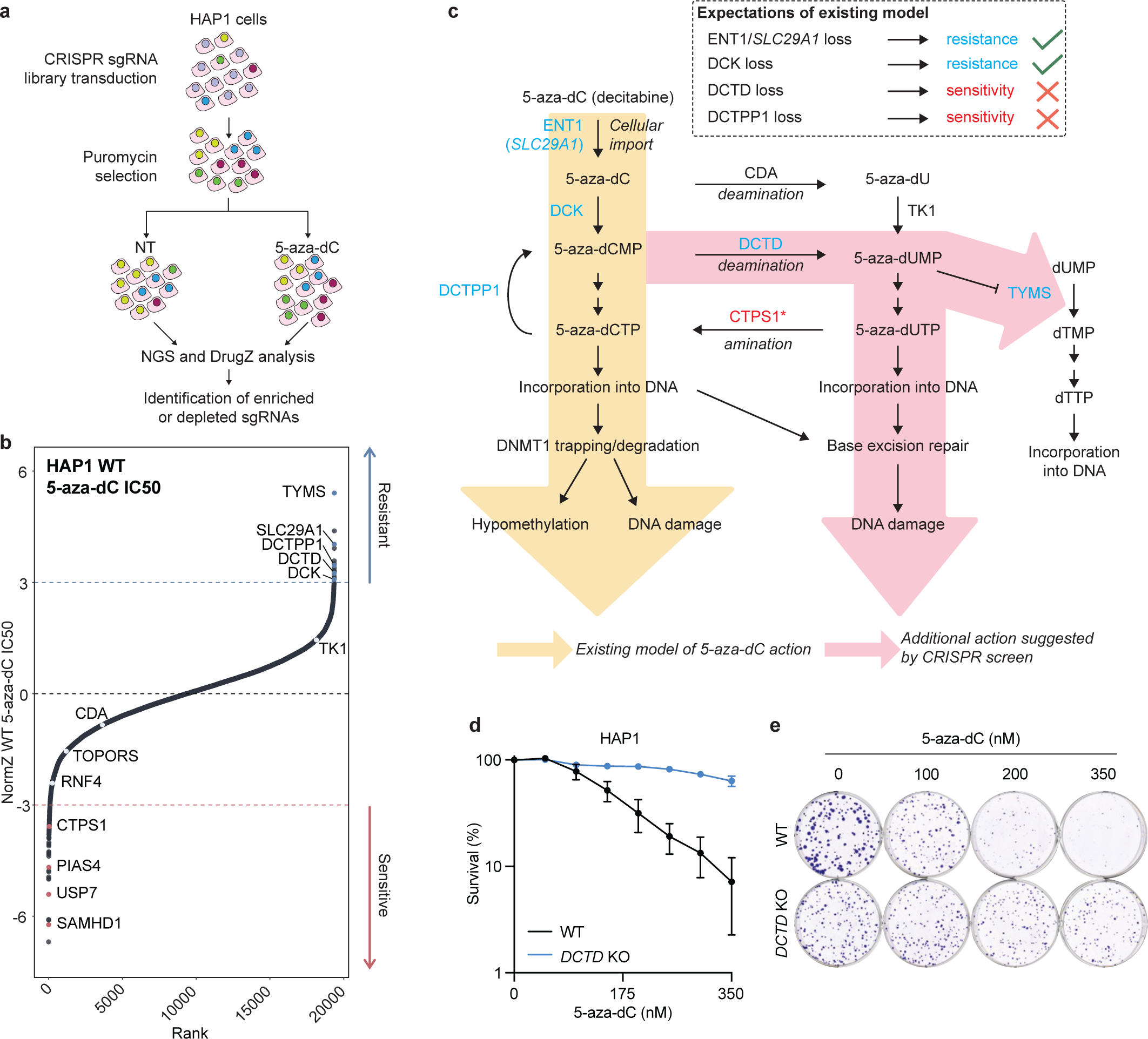
Loss of DCTD confers resistance to 5-aza-dC. (a) Schematic outlining genome-wide CRISPR/Cas9 screen with 5-aza-dC in HAP1 cells. (b) Rank plot displaying selected hits from CRISPR/Cas9 screen with 5-aza-dC, outlined in (a); dotted lines at NormZ scores of +3/-3 indicate threshold for resistance/sensitivity hits, respectively. (c) Schematic detailing the existing model of 5-aza-dC action and the additional action suggested by our CRISPR screen. Factors whose loss confers resistance/sensitivity to 5-aza-dC in the CRISPR screen in (b) are displayed in blue/red, respectively. * denotes inferred enzymatic activity based on its yeast homologue. (d) Clonogenic survival assays in WT and *DCTD* KO HAP1 cells treated with 5-aza-dC and stained 6 days later; n = 3 biological replicates, error bars ± SEM. (e) Representative images from (d) of cells at selected 5-aza-dC doses.

Strikingly, and challenging the prevailing model of the mechanism of decitabine action, inactivation of *DCTD* or *DCTPP1* conferred 5-aza-dC resistance in our CRISPR screen (Fig 1b-c). DCTD deaminates dCMP/5-aza-dCMP to dUMP/5-aza-dUMP^44^, while DCTPP1 dephosphorylates dCTP to dCMP (and presumably 5-aza-dCTP to 5-aza-dCMP), thus regenerating a substrate for DCTD (Fig 1c). To validate this, we assessed the 5-aza-dC sensitivity of *DCTD* knockout (KO) HAP1 cells in clonogenic survival assays and observed profound 5-aza-dC resistance compared to wild-type (WT) cells (Fig 1d-e). 5-aza-dUMP, generated by DCTD’s action on 5-aza-dCMP, has been demonstrated to bind TYMS^44^, although the functional consequence of this interaction has been unclear. Notably, *TYMS* inactivation also conferred strong 5-aza-dC resistance in our CRISPR screen (Fig 1b), possibly suggesting that the interaction between 5-aza-dUMP and TYMS contributes to cytotoxicity. Together, these findings highlight a mechanism of 5-aza-dC cytotoxicity that occurs through generation of 5-aza-dUMP.

### 5-aza-dC deamination drives DNMT1-independent cytotoxicity

Next, we wanted to determine whether the unexpected 5-aza-dC resistance upon loss of DCTD was related to differences in DNMT1-DPC formation. Therefore, we used the recently developed Purification of x-linked Proteins (PxP) assay^38^ (Fig 2a) to monitor induction of DNMT1-DPCs by 5-aza-dC in *DCK* and *DCTD* KO cells. In WT cells, DNMT1-DPCs were induced within 3 hours of 5-aza-dC treatment (Fig 2b). By 6 hours of treatment, the level of DNMT1-DPCs was reduced, presumably reflecting the progressive degradation of DNMT1-DPCs. Consistently, total cellular DNMT1 was concomitantly depleted, as evident from analysis of input samples, reminiscent of previous observations^38,45^ (Fig 2b). In *DCK* KO cells however, 5-aza-dC is not expected to be incorporated into DNA because DCK is required for the first phosphorylation step in the generation of 5-aza-dCTP. In agreement with this, we only detected minimal 5-aza-dC-induced DNMT1-DPCs in *DCK* KO cells and only at the later time point of 6 hours (Fig 2b). It is possible that the residual DNMT1-DPCs detected in these circumstances are caused by 5-aza-dCTP incorporation arising downstream of deamination of 5-aza-dC to 5-aza-dU by CDA, followed by triphosphorylation and conversion of 5-aza-dUTP to 5-aza-dCTP by CTPS1. In line with this idea, the *Saccharomyces cerevisiae* homologue of CTPS1 has been described to convert dUTP to dCTP^46,47^(Supplemental Fig 1). In contrast, to our findings with *DCK* KO cells, DNMT1-DPC induction in *DCTD* KO cells was similar to that in WT cells, (Fig 2b), suggesting that resistance of *DCTD* KO cells to 5-aza-dC is not mediated by reduced formation of DNMT1-DPCs. These data thus demonstrate that the main cytotoxic action of 5-aza-dC, upon its deamination and production of 5-aza-dUMP, is DPC-independent.

**Figure 2.**
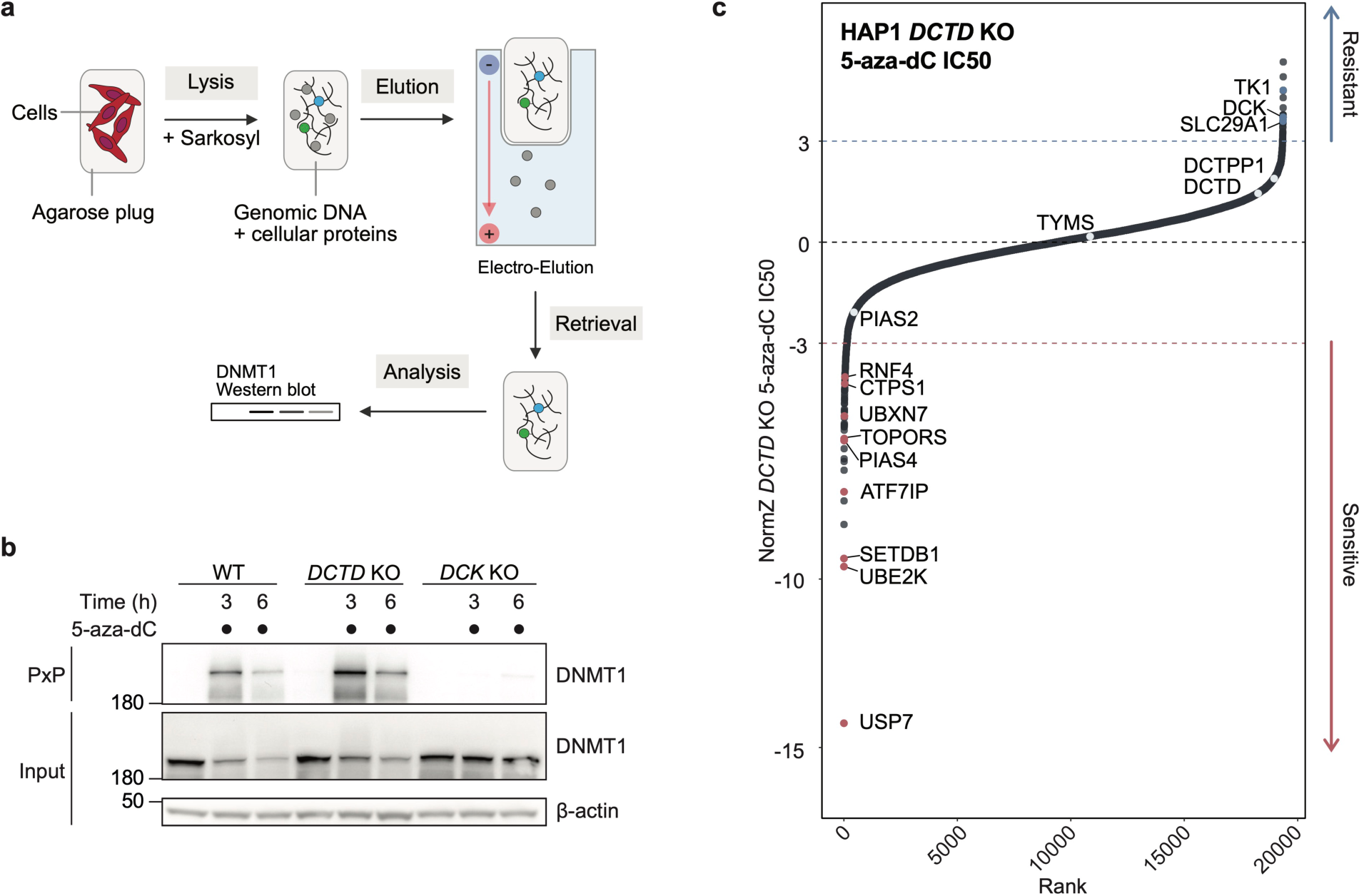
5-aza-dC cytotoxicity is driven by DNMT1-dependent and -independent mechanisms. (a) Schematic detailing the PxP assay. Cells are harvested and cast into low-melt agarose plugs. Plugs are transferred to denaturing lysis buffer. After lysis is completed, plugs are placed into the pockets of a SDS-PAGE-gel and non-crosslinked proteins are eluted by electrophoresis. Finally, plugs are retrieved from the gel pockets and boiled with LDS sample buffer. For DPC detection, samples are run on SDS-PAGE gels and quantified by western blotting. (b) DNMT1-DPC formation assessed by PxP in WT, *DCTD* KO and *DCK* KO HAP1 cells treated with 5-aza-dC (1 μM) for the indicated times. (c) Rank plot of a genome-wide CRISPR/Cas9 screen in *DCTD* KO HAP1 cells treated with 5-aza-dC; dotted lines at NormZ scores of +3/-3 indicate threshold for resistance/sensitivity hits, respectively.

To gain further insight into mechanisms of 5-aza-dC resistance, we performed a genome-scale CRISPR screen in *DCTD* KO cells. Strikingly, in contrast to our 5-aza-dC screen in WT cells (Fig 1b), in *DCTD* KO cells, gRNAs targeting *TYMS*, *DCTD* or *DCTPP1* no longer conferred 5-aza-dC resistance, while gRNAs against *DCK* still conferred strong resistance (Fig 2c). In addition, we identified several factors whose loss caused strong 5-aza-dC sensitivity (Fig 2c). Most prominently, we identified factors which have established roles in replication-independent repair of DPCs^36,37^, and the deubiquitylating enzyme USP7, which regulates the DPC protease SPRTN^48^ among various other processes^49^. We additionally identified factors with unclear roles in 5-aza-dC tolerance such as the dual SUMO1 and ubiquitin E3 ligase TOPORS and the ubiquitin E2 conjugating enzyme UBE2K (Fig 2c). This indicates that in the absence of 5-aza-dC deamination, the relative contribution of DNMT1-DPCs to 5-aza-dC-induced cytotoxicity substantially increases.

### TOPORS is recruited to DNMT1-DPCs and promotes DPC tolerance

In light of the above findings, we speculated that in addition to identifying known DPC repair factors such as RNF4 and PIAS4, our CRISPR screen for 5-aza-dC sensitivity in *DCTD* KO cells may have uncovered as-yet unrecognised DPC repair factors. To explore this, we developed a variation of iPOND (isolation of proteins on nascent DNA)^50^, which we term iPOND-DPC, enabling determination of the proximal proteome of DNMT1-DPCs. Briefly, HeLa-TREx cells were synchronised via double thymidine block and released into early/mid S-phase. Cells were co-treated with 5-ethynyl-deoxyuridine (EdU) and 5-aza-dC for 30 minutes to ensure their co-incorporation into nascent DNA, allowing the subsequent specific isolation of DPC-containing chromatin (Fig 3a). To this end, cells were crosslinked with 1% formaldehyde followed by biotinylation of EdU by using a click reaction with biotin-azide, DNA shearing and streptavidin-bead mediated retrieval of DPC-containing DNA fragments. To validate our experimental protocol, we analysed the flowthrough and iPOND-DPC samples by Western blotting with antibodies for DNMT1, histone H3 and tubulin (Fig 3b). Histone H3, but not tubulin was retrieved on streptavidin beads when cells were treated with EdU, indicating successful purification of nascent chromatin. Similarly, DNMT1 was detected on nascent chromatin, consistent with its key role in maintenance of DNA methylation. Importantly, when cells were additionally treated with 5-aza-dC, the DNMT1 signal increased in iPOND-DPC samples, while histone H3 levels remained unchanged, indicating the formation of persistent DNMT1-DPC crosslinks in post-replicative chromatin (Fig 3b).

**Figure 3.**
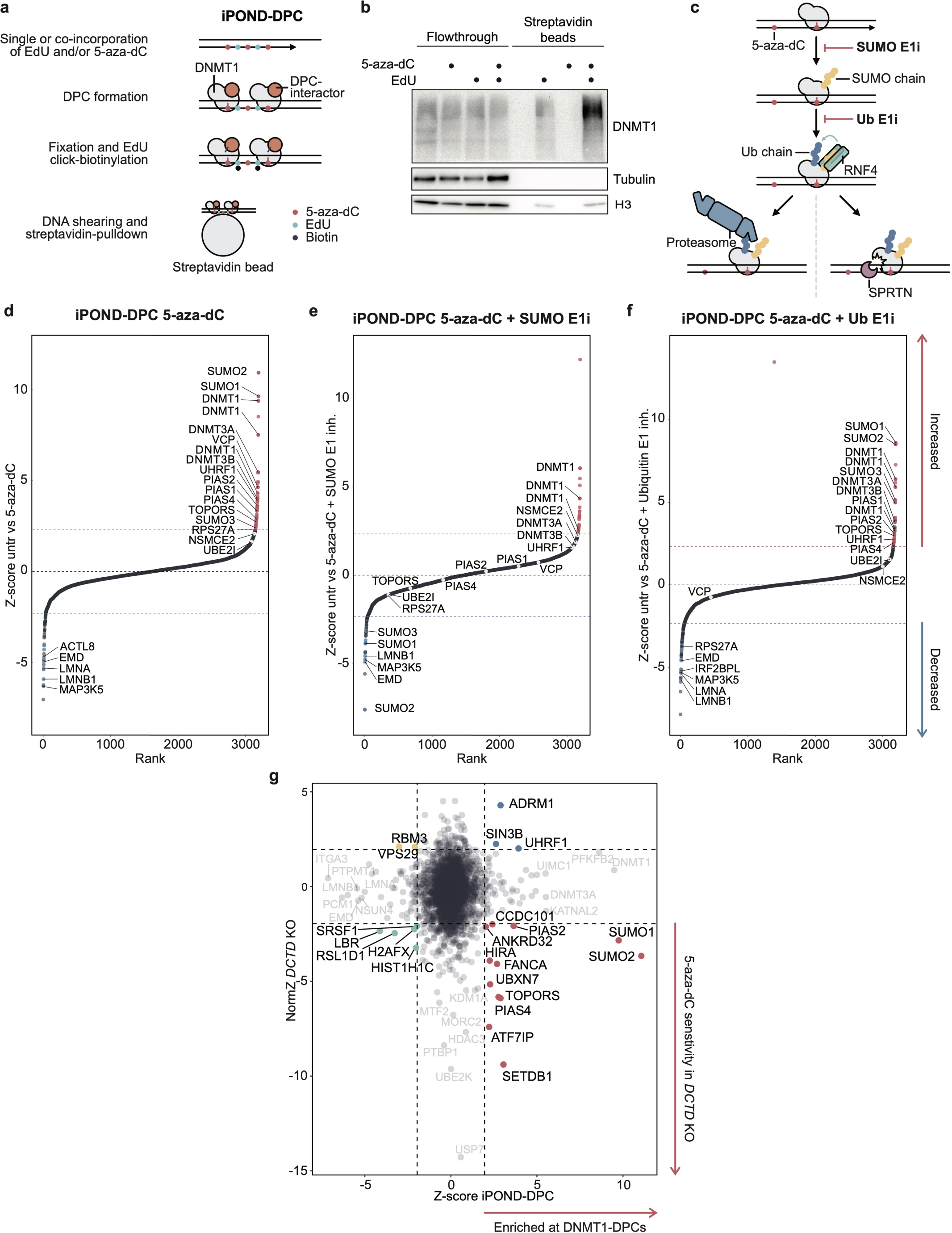
iPOND-DPC identifies SUMO- and ubiquitin-dependent DNMT1-DPC-proximal factors. (a) Schematic outlining the iPOND-DPC approach. (b) HeLa TREx cells were treated with either 5-aza-dC (10 μM), EdU (10 μM) or both and processed as depicted in (a). Samples were analysed by Western Blot using the indicated antibodies. (c) Schematic outlining global-genome (GG-) DPC repair and the impacts of SUMO or ubiquitin E1 inhibitors on this pathway. (d) Ranked standardised enrichment of proteins detected by iPOND-DPC/MS from 5-aza-dC treated over untreated cells. Dotted lines indicate a threshold of ± 2.326. Four replicates were measured. (e) Ranked standardised enrichment of proteins detected by iPOND-DPC/MS from co-treatment with 5-aza-dC and SUMO E1i over untreated cells. Dotted lines indicate a threshold of ± 2 2.326. Four replicates were measured. (f) Ranked standardised enrichment of proteins detected by iPOND-DPC/MS from co-treatment with 5-aza-dC and TAK-243 over untreated cells. Dotted lines indicate a threshold of ± 2.326. Four replicates were measured. (g) Scatter plot comparing standardised enrichment scores of iPOND-DPC/MS from (d) with NormZ score of the top 10% of hits from our 5-aza-dC CRISPR/Cas9 screen in *DCTD* KO cells from Fig 2c. Dotted lines indicate a threshold of ± 2.326.

Next, we set out to determine the proximal proteome of DNMT1-DPCs using liquid chromatography with tandem mass tag (TMT)-multiplexed mass spectrometry. To recapitulate the successive stages of SUMO and ubiquitin-dependent DNMT1-DPC repair ^35–38^(Fig 3c), we compared proteins identified in iPOND-DPC samples of untreated cells with those of cells treated with 5-aza-dC, co-treated with both 5-aza-dC and the SUMO E1 inhibitor (SUMO E1i) ML-792, or co-treated with 5-aza-dC and the ubiquitin E1 inhibitor (Ub E1i) TAK-243. Upon 5-aza-dC treatment, DNMT1, DNMT3A and DNMT3B, UHRF1, SUMO2 and SUMO1 were among the most enriched proteins in iPOND-DPC samples (Fig 3d). We also identified factors previously shown to be involved in DPC repair such as the SUMO E3 ligase PIAS4^36,37^, VCP/p97^38,51,52^ and most proteasome subunits (Fig 3d, Supplemental Fig 2a; no peptides for RNF4 were detected in any of our samples). Inhibition of SUMOylation abrogated the recruitment of proteins involved in SUMO conjugation, including PIAS1-4, UBE2I and TOPORS as well as VCP/p97 and to some extent the proteasome, but retained DNMT1 enrichment (Fig 3e, Supplemental Fig 2b). In contrast, while inhibition of ubiquitylation completely abrogated the recruitment of VCP/p97 and proteasome subunits, it did not diminish recruitment of PIAS1-4 or TOPORS (Fig 3f, Supplemental Fig 2c-d).

To uncover proteins that are recruited to DNMT1-DPCs and whose loss causes 5-aza-dC sensitivity, we compared proteins identified to be in proximity of DNMT1-DPCs by iPOND-DPC with the top 2.5% of sensitivity hits in our 5-aza-dC CRISPR screen in *DCTD* KO cells after plotting the scores of all mutually-occurring hits. This analysis identified the histone methyltransferase SETDB1 and its associated factor ATF7IP, the SUMO ligases PIAS2 and PIAS4, the VCP/p97 adaptor UBXN7 and the interstrand DNA crosslink (ICL) repair protein FANCA (Fig 3g, lower right quadrant). Additionally, this analysis highlighted the dual ubiquitin and SUMO1 E3 ligase TOPORS^53–56^. Furthermore, we noted that our iPOND-DPC studies indicated that TOPORS is recruited to DNMT1-DPCs in a SUMO-dependent but ubiquitin-independent manner (Fig 3c,e-f, Supplemental Fig 2d). Together, these findings, coupled with results from our genetic screen in *DCTD* KO cells, pointed to a direct and important role for TOPORS in response to 5-aza-dC-induced DNMT1-DPCs.

### TOPORS acts downstream of DNMT1-DPC SUMOylation to counter 5-aza-dC toxicity

To begin to explore TOPORS functions, we established *TOPORS* KO clones of human *TP53* KO RPE1 cells. These cells displayed hypersensitivity towards 5-aza-dC when compared to isogenic controls (Fig 4a; Supplemental Fig 3a), a phenotype that we also observed in commercially-available *TOPORS* KO HAP1 cells (Fig 4b). Furthermore, siRNA-mediated depletion of TOPORS also caused 5-aza-dC hypersensitivity in human U2OS cells (Fig 4c; Supplemental Fig 3b). In addition, we profiled *TOPORS* HAP1 cells for other hypersensitivity phenotypes relevant to genome stability, and observed hypersensitivity towards formaldehyde and camptothecin, which induce DPCs, but not to ionising radiation, which causes DNA breaks (Fig 4d-f).

**Figure 4.**
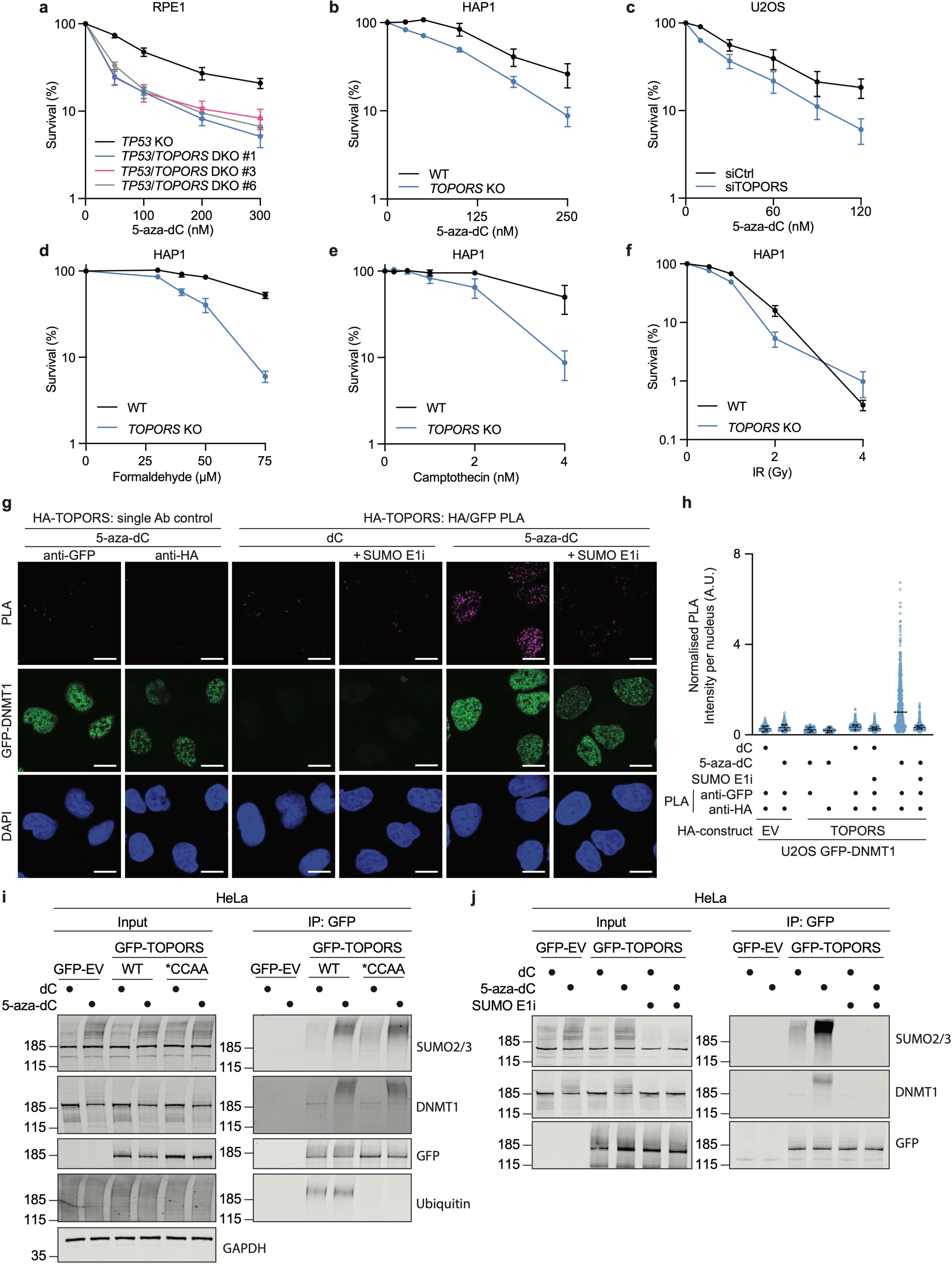
TOPORS is a SUMO-dependent DPC tolerance factor. (a) Clonogenic survival assays in *TP53* KO and three clonally-derived *TP53*/*TOPORS* DKO RPE1 cell lines treated with 5-aza-dC and stained 6 days later; n = 4 biological replicates, error bars ± SEM. (b) Clonogenic survival assays in WT and *TOPORS* KO HAP1 cells treated with 5-aza-dC and stained 6 days later; n = 4 biological replicates, error bars ± SEM. (c) Clonogenic survival assays in WT U2OS cells transfected with siCtrl or siTOPORS and treated with 5-aza-dC and stained 6 days later; n = 3 biological replicates, error bars ± SEM. (d-f) Clonogenic survival assays in WT and *TOPORS* KO HAP1 cells treated with formaldehyde (d), camptothecin (e) and ionizing radiation (IR; e) and stained 6 days later; n = 3 biological replicates, error bars ± SEM. (g) Proximity ligation assay in U2OS cells expressing GFP-DNMT1 and HA-TOPORS, released from a single thymidine block and treated with deoxycytidine (dC), 5-aza-dC and/or SUMO E1i for 1 hour before pre-extraction of non-chromatin-bound proteins and fixation. (h) Quantification of per-nucleus mean PLA intensities (blue dots) from (g), normalised to the median from the 5-aza-dC-treated U2OS GFP-DNMT1/HA-TOPORS condition. Black dots display the median normalised PLA intensity of each biological replicate for each condition; n = 4 independent biological replicates error bars ± SEM. (i) Co-immunoprecipitation of GFP from extracts of HeLa cells expressing GFP (EV), GFP-TOPORS^WT^ or GFP-TOPORS^CCAA^, released from thymidine block into S-phase and treated with dC or 5-aza-dC for 1 hour, followed by western blotting for indicated proteins. (j) Co-immunoprecipitation of GFP from extracts of HeLa cells expressing GFP (EV) or GFP-TOPORS^WT^, released from thymidine block into S-phase and treated with dC, 5-aza-dC and/or SUMO E1i for 1h, followed by western blotting for indicated proteins.

To build on these findings, and confirm our iPOND-DPC datasets, we established cell lines stably expressing HA-tagged TOPORS or the empty vector (EV) in U2OS cells also constitutively expressing green-fluorescent protein (GFP)-tagged DNMT1 (U2OS GFP-DNMT1). We synchronised these cells in S-phase by thymidine block and observed recruitment of TOPORS to DNMT1 upon release into 5-aza-dC treatment, as assessed by Proximity Ligation Assay (PLA) (Fig 4g-h; see Supplemental Fig 4a for representative images in U2OS GFP-DNMT1 HA-EV cells). In agreement with our previous observations, co-treatment of 5-aza-dC with SUMO E1i returned the PLA signal to background levels, indicating abolition of TOPORS recruitment to DNMT1-DPCs (Fig 4g-h). TOPORS is a known SUMO interactor and contains five SUMO-interaction motifs (SIMs)^57^, suggesting that SUMO-dependent recruitment of TOPORS to DNMT1-DPCs could be mediated by direct interactions with SUMO chains formed on DPCs. Consistent with this idea, immunoprecipitation of WT GFP-TOPORS or a ubiquitylation-defective GFP-TOPORS RING domain mutant (GFP-TOPORS^CCAA^) from 5-aza-dC-treated HeLa cells revealed robust interactions between GFP-TOPORS and high molecular weight SUMOylated proteins, when compared to the deoxycytidine treatment (Fig 4i). Moreover, an interaction between GFP-TOPORS and heavily modified DNMT1 was strongly induced by 5-aza-dC treatment, likely corresponding to extensively SUMOylated and ubiquitylated 5-aza-dC-induced DNMT1-DPCs^35^ (Fig 4i). While we detected an interaction between WT GFP-TOPORS and a high molecular weight ubiquitylated interactor, this was not overtly 5-aza-dC-inducible but undetectable in co-immunoprecipitates of GFP-TOPORS^CCAA^ (Fig 4i). Notably, and in agreement with our iPOND-DPC and PLA data, cotreatment of GFP-TOPORS-expressing HeLa cells with SUMO E1i alongside 5-aza-dC essentially abrogated GFP-TOPORS’ interactions with SUMOylated proteins and DNMT1 (Fig 4j). Collectively, these data corroborate a direct role for TOPORS in response to 5-aza-dC-induced DNMT1-DPCs that entails its SUMO-dependent recruitment to DPCs.

### TOPORS acts in parallel to RNF4 and UBE2K in mediating DNMT1-DPC tolerance

To better understand the role of TOPORS in promoting cellular tolerance to DNMT1-DPCs and thus towards 5-aza-dC, we performed a further 5-aza-dC CRISPR screen in *TOPORS* KO HAP1 cells. This revealed that *TOPORS* KO cells are strongly sensitised to 5-aza-dC by additional loss of PIAS4, RNF4 or the ubiquitin-conjugating E2 enzyme UBE2K (Fig 5a). Our CRISPR screen thus suggested that TOPORS acts independently of the PIAS4-RNF4 axis for DNMT1-DPC repair. Indeed, siRNA-mediated depletion of RNF4 caused dramatic sensitization of *TOPORS* KO HAP1 cells to 5-aza-dC (Fig 5b; Supplemental Fig 4b), while siTOPORS caused profound additional sensitivity in *RNF4* KO HeLa cells (Fig 5c). Furthermore, cellular fitness, measured by proliferation rate, was severely compromised by the loss of both TOPORS and RNF4 in HeLa cells (Fig 5d-e) and we were unable to knock out *TOPORS* in *RNF4* KO HeLa cells, possibly reflecting an inability of TOPORS- and RNF4-deficient cells to tolerate endogenously arising DPCs.

**Figure 5.**
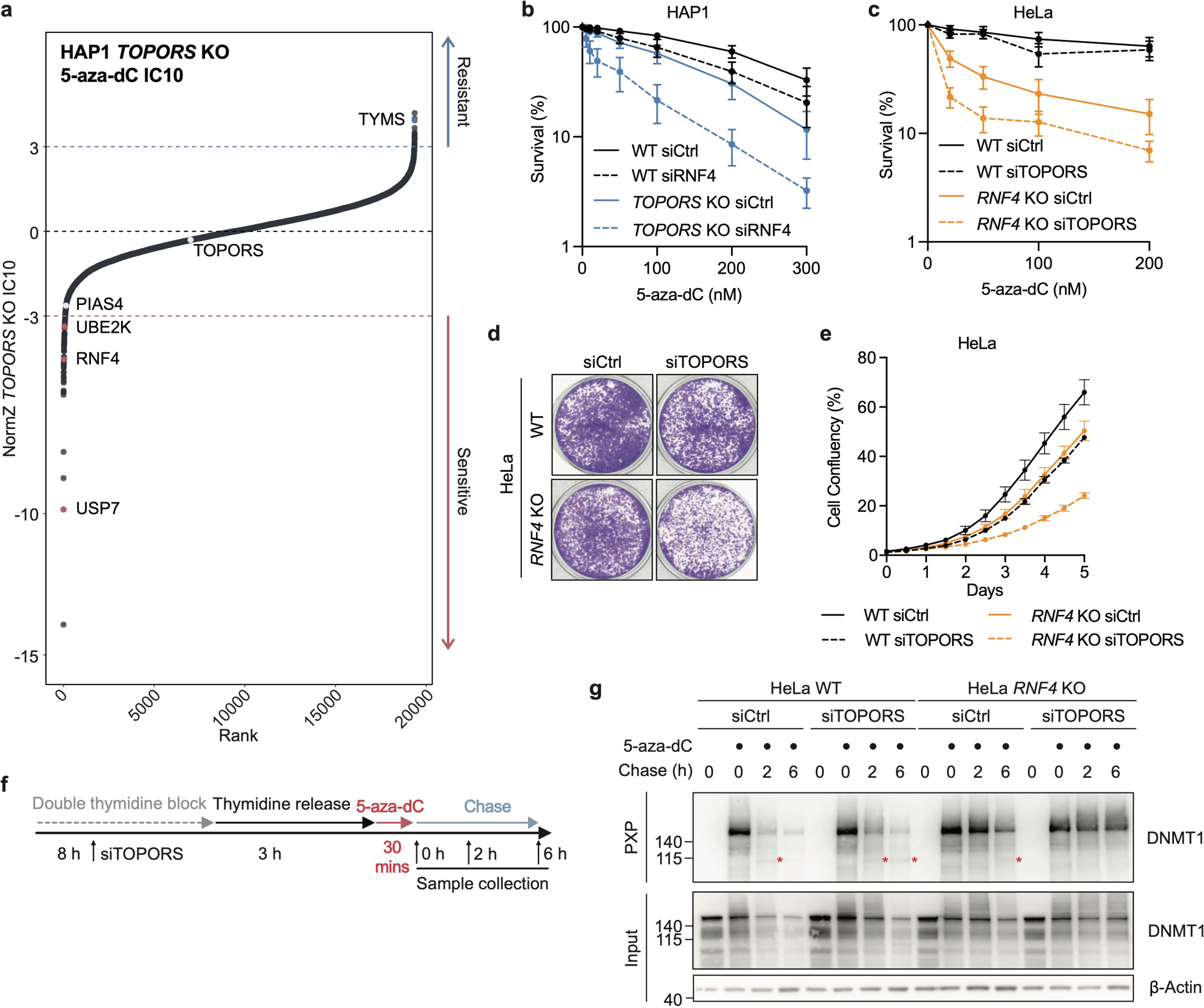
TOPORS and RNF4 operate in parallel to promote DPC degradation. (a) Rank plot of a genome-wide CRISPR/Cas9 screen in *TOPORS* KO HAP1 cells treated with 5-aza-dC; dotted lines at NormZ scores of +3/-3 indicate threshold for resistance/sensitivity hits, respectively. (b) Clonogenic survival assays in WT and *TOPORS* KO HAP1 cells transfected with siCtrl or siRNF4, treated with 5-aza-dC 72h after transfection and stained 6 days later; n=4 biological replicates, error bars ± SEM. (c) Clonogenic survival assays in WT and *RNF4* KO HeLa cells transfected with siCtrl or siTOPORS, treated with 5-aza-dC 72h after transfection and stained 6 days later; n=4 biological replicates, error bars ± SEM. (d) Representative image of cell confluency of WT and *RNF4* KO HeLa cells transfected with siCtrl or siTOPORS. (e) Cell confluency of WT and *RNF4* KO HeLa cells transfected with siCtrl or siTOPORS for a period of 5 days; n=2 biological replicates. Cell confluency was monitored using IncuCyte live cell imaging. n=2 biological replicates. (f) Treatment schematic for PxP assay in (g). (g) HeLa WT or *RNF4* KO were transfected with the indicated siRNAs, synchronised by double thymidine block and treated with 5-aza-dC (10 μM) as depicted in (f). DNMT1-DPC were isolated using PxP and analysed by western blotting using the indicated antibodies. Asterisks (*) indicate SPRTN dependent DNMT1 cleavage fragment.

Interestingly, our CRISPR screen in *TOPORS* KO cells indicated that inactivation of the ubiquitin E2 conjugating enzyme UBE2K also caused further sensitisation to 5-aza-dC (Fig 5a). Indeed, we found that UBE2K loss caused 5-aza-dC hypersensitivity that was dramatically exacerbated by additional loss of TOPORS (Supplemental Fig 4c-e). To explore the relationship between UBE2K and RNF4, we depleted RNF4 using siRNA in *UBE2K* KO cells, which caused increased 5-aza-dC hypersensitivity (Supplemental Fig 4f). Furthermore, RNF4 depletion further sensitised *UBE2K*/*TOPORS* double knockout (DKO) cells to 5-aza-dC (Supp Fig 4g). These data indicated that UBE2K plays a role in 5-aza-dC tolerance that is independent of both TOPORS and RNF4. In support of this, despite being detected in the vicinity of nascent DNA in our iPOND-DPC experiments, UBE2K - in contrast to TOPORS - was not further enriched upon DNMT1-DPC induction by 5-aza-dC. These findings place TOPORS in a DPC tolerance pathway acting alongside UBE2K and the RNF4 axis and suggest that these two E3 ubiquitin ligases perform at least partially overlapping functions necessary for survival when cells are challenged with 5-aza-dC.

While proteasomal degradation and SPRTN-dependent cleavage of DNMT1-DPCs is severely delayed in *RNF4* KO cells, it is not entirely abrogated^38^. Given that the residual DNMT1-DPC proteolysis also depends on SUMOylation and ubiquitylation^38^, we hypothesised that TOPORS promotes polyubiquitylation and degradation of DNMT1-DPCs in parallel to RNF4. TOPORS’ implication in the repair of endogenous DPCs could explain the proliferation defect observed in cells depleted of both RNF4 and TOPORS. We employed the PxP assay^38^ (Fig 2a) to examine the resolution of DNMT1-DPCs in WT and *RNF4* KO HeLa cells upon TOPORS depletion by siRNA. Therefore, we synchronised cells in early/mid-S-phase before DNMT1-DPCs’ induction with a 30-minute 5-aza-dC pulse (Fig 5f). Following treatment, cells were either harvested directly or allowed to recover in drug-free medium for two or six hours. As shown previously^38^, DNMT1-DPCs were readily detectable following 5-aza-dC treatment but were swiftly lost in WT cells following wash-out of the drug (Fig 5g; a proteolytic fragment arising from SPRTN-dependent cleavage of the DNMT1-DPC is highlighted with an asterisk). In *RNF4* KO HeLa cells, DNMT1-DPC degradation was delayed and only observable at the six-hour time point. Strikingly, while siRNA-mediated TOPORS depletion in WT cells caused only a modest impairment in DNMT1-DPC degradation, TOPORS loss in *RNF4* KO cells virtually abolished DNMT1-DPC degradation at the time points tested (Fig 5g). Additionally, corresponding deficiencies in DNMT1 degradation across these conditions were also visible in the PxP ‘input’ samples (Fig 5g).

Together, our results support a model in which, in parallel with RNF4 recruitment, TOPORS is recruited to DNMT1-DPCs following their SUMOylation, where it promotes DPC polyubiquitylation and subsequent proteolysis to promote cell viability upon decitabine/5-aza-dC treatment.

## Discussion

In this study, we explored the cytotoxic effects of the hypomethylating agent 5-aza-dC by using unbiased genetic screens. We combined these insights with the first mapping of the proximal proteome of 5-aza-dC-induced DNMT1-DPCs. Our results shed light on two important modes of 5-aza-dC action. First, we reveal that a substantial amount of cytotoxicity originates from deamination of 5-aza-dC by DCTD and the generation of 5-aza-dUMP, highlighting DCTD loss as an important mechanism of decitabine resistance. Second, we find that the dual SUMO1/ubiquitin E3 ligase, TOPORS, is a key player in global-genome DPC repair that promotes the degradation of 5-aza-dC-induced DNMT1-DPCs.

Global-genome DPC repair is initiated upon SUMOylation of the DPC^35–37^. Our data demonstrate that ensuing SUMO-dependent ubiquitylation is not only promoted by RNF4 but also by TOPORS. Indeed, loss of RNF4 compromises - but does not abolish - the SUMO-dependent degradation of DNMT1-DPCs^37,38^, which is only blocked upon loss of both TOPORS and RNF4, rendering cells extremely sensitive to 5-aza-dC. Like RNF4, TOPORS is recruited to SUMOylated DNMT1-DPCs and the simultaneous depletion of both TOPORS and RNF4 causes a strong fitness defect. Therefore, TOPORS appears to act in parallel to RNF4 to promote global-genome DPC repair with some level of mutual redundancy. However, this redundancy is likely limited. While RNF4 loss is embryonic lethal in mice^59^, mutations in TOPORS are associated with a variant of retinitis pigmentosa^60^, characterised by apoptotic rod cells, and in Joubert syndrome^61^, a rare disease characterised by brain stem anomalies and in some cases retinal dystrophy. In fruit flies, TOPORS was observed to be localised to dedicated nuclear compartments where it regulates the activity of chromatin insulator complexes^62^. This implies that TOPORS and RNF4 have substantial non-overlapping functions *in vivo*. While subsequent studies will hopefully further explore and define the relationship between the two enzymes, it is conceivable that DNMT1-DPCs might be preferentially ubiquitylated by TOPORS or RNF4 depending on the chromatin context but can be modified by either if the lesion persists for a longer period of time. TOPORS is an unusual E3 ligase in that it promotes conjugation of both ubiquitin conjugation via its N-terminal RING domain, and SUMO1 conjugation via a region in its unstructured C-terminal tail. Our findings demonstrate that TOPORS is recruited to DNMT1-DPCs in a SUMO-dependent manner. However, whether SUMOylation is only required to recruit TOPORS or is also activating the enzyme as in the case of RNF4 remains unclear^63^. In addition, understanding whether TOPORS’ role in DPC repair requires its own SUMO E3 ligase activity or relies on other SUMOylating enzymes is an important future goal.

The formation of DNMT1-DPCs is toxic upon loss of global genome DPC repair factors. However, our genetic screens identified an additional DNMT1-independent mode of action that is dominant in WT cells and originates from 5-aza-dCMP deamination to 5-aza-dUMP by DCTD. The generation of 5-aza-dUMP through deamination by DCTD has been shown previously^22,23,44^, although to our knowledge we provide the first genetic evidence that DCTD underlies 5-aza-dC cytotoxicity through 5-aza-dUMP production. Notably, our CRISPR/Cas9 screens indicate that even the residual 5-aza-dC sensitivity in *DCTD* KO cells appears to stem in part from 5-aza-dUMP generation. In *DCTD* KO but not WT cells, *TK1* loss causes 5-aza-dC resistance. This might suggest that in the absence of DCTD, the sequential action of CDA and TK1 on 5-aza-dC can generate 5-aza-dUMP and impart cytotoxicity. The relative genetic dependencies of WT and *DCTD* KO cells upon 5-aza-dC treatment raises interesting questions about the mechanism of 5-aza-dUMP’s cytotoxicity. We envision two possible scenarios. First, given that 5-aza-dUMP interacts with TYMS *in vitro* and in cells^44,64^, it could therefore inhibit the action of TYMS, as many other chemotherapeutics do^65,66^. TYMS inhibition may be caused by the nitrogen substitution in place of a carbon at the 5’ position within the pyrimidine ring of 5-aza-dUMP, which is expected to block its methylation by TYMS. The fact that in our CRISPR/Cas9 screens, loss of TYMS caused 5-aza-dC resistance in WT cells but had no effect in *DCTD* KO cells provides support for this scenario. Second, incorporation of 5-aza-dUTP into DNA could activate base excision repair (BER) and result in persistent DNA single strand break (SSB) formation, which could be exacerbated by simultaneous inhibition of TYMS. Indeed, inhibition of the SSB repair factor PARP1 is known to synergise with 5-aza-dC^67,68^. However, while SSB induction by 5-aza-dC has been reported^67,69^, it remains unclear whether these SSBs arise as a result of BER action on genomic 5-aza-dCTP, 5-aza-dUTP, dUTP or all of these. The relative contributions of these non-mutually exclusive scenarios is an exciting issue for future exploration.

Taken together, our findings have important implications for the understanding and clinical applications of 5-aza-dC, and might serve as a starting point for the development of new candidate biomarkers to guide patient stratification. Our results suggest that in the presence of proficient DPC repair, much of 5-aza-dC’s cytotoxic effect originates not from DNMT1-DPCs but from the generation of 5-aza-dUMP. Given that hypomethylation is driven by DNMT1-DPC degradation, our work suggests that at clinically relevant doses, the origins of cytotoxicity are independent of DNA hypomethylation. This insight might help guide the development of new strategies to maximise either hypomethylation or cytotoxic effects depending on the clinical goal. Importantly, our data also highlights that the accepted mode of 5-aza-dC detoxification through deamination must be revisited.

### Cell Culture

Cell lines used in this study were maintained as detailed in Supplementary Table 1.

TOPORS was knocked out in*TP53* KO RPE1 cells stably expressing Cas9^70^ by transient transfection of sgRNA duplexes of an Alt-R tracrRNA (IDT) and a TOPORS-specific Alt-R crRNA (Supplementary Table 2), formed as per the manufacturer’s recommendations and transfected using TransIT-LT1 (Mirus) as per the manufacturer’s recommendations. 48h later, cells were seeded at >1 cell/well in 96-well plates and expanded as clonal populations. To knock out *TOPORS* in *UBE2K* KO HAP1 cells, ribonucleoprotein (RNP) complexes were prepared between spCas9 (IDT) and the Alt-R sgRNA duplex prepared as described previously according to IDT’s Alt-R protocol. RNPs were electroporated into HAP1 cells using a Neon NxT Electroporation System (ThermoFisher) as per the manufacturer’s instructions. To validate candidate *TP53*/*TOPORS* DKO (double KO) and *UBE2K*/*TOPORS* DKO clones, genomic DNA extracts were prepared using the DNeasy Blood & Tissue Kit (Qiagen) as per the manufacturer’s instructions. Clones were validated by TIDE analysis^71^ following Sanger sequencing of PCR amplicons containing the targeted region of *TOPORS* using primers detailed in Supplementary Table 2.

To generate lentiviruses, HEK293T LentiX cells were transfected with TransIT-LT1 (Mirus) transfection reagent as well as a construct of interest (Supplementary Table 2) and the psPAX2 (Addgene 12260), and pMD2.G (Addgene 12259) constructs, according to the manufacturer’s protocol. 48 hours after transfection, the viral supernatant was collected and filtered through a 0.45μm sterile Millex-GP filter unit (Merck). The lentivirus was stored at −80°C until use.

U2OS cells stably expressing GFP-DNMT1 (U2OS GFP-DNMT1) were established by transfecting cells with pEGFP-DNMT1 using TransIT-LT1 (Mirus) as per the manufacturer’s instructions, then selecting transfected cells with 1mg/ml G418 (Gibco) for 7 days. G418-resistant cells were then seeded into 96-well plates at a concentration of 0.5 cells/well and after two weeks monoclonal cell lines were validated by GFP fluorescence and immunoblotting. U2OS GFP-DNMT1 cells were then infected with lentivirus-containing supernatants with pHA-EV-lentipuro or pHA-TOPORS-lentipuro and 48h later selected with 2μg/ml puromycin for 48h. Puromycin-resistant polyclonal cell populations were then validated by immunofluorescence against the HA-tag.

### CRISPR/Cas9 screens

2.5×108 WT HAP1, *DCTD* KO and *TOPORS* KO HAP1 cells were infected with the pre-packaged genome-wide all-in-one Brunello lentiviral library (Addgene 73179) at a multiplicity of infection (MOI) of 0.2. Following lentiviral integration, the transfected cells were selected with puromycin (0.6μg/ml). Puromycin-resistant cells were passaged and expanded as required for 8 days. During this period, the cells were split into two independent replicates at a library coverage of 500x, that is, a total of 50 ×10^6^ cells per condition. After cell expansion, each replicate was treated with decitabine (30nM for WT cells, 250nM for *DCTD* KO cells and 12nM for *TOPORS* KO cells) or DMSO, for a period of ten days. The first day of the ten-day treatment is Day 0 (T0) of the screen. During the course of the screen, cells were passaged and re-treated every other day, at a coverage of 500x.

For each replicate and treatment condition, cell pellets were collected on the first (T0) and last (T10) days of the screen, at a coverage of 500X. Upon harvesting of the cell pellets, genomic DNA was extracted using the QIAamp Blood Maxi kit (QIAGEN). Following the precipitation and purification of DNA in 70% ethanol, the DNA was dissolved to a concentration of 500ng/μl in H_2_O. The sgRNA sequences of each DNA sample were then amplified by PCR using the NEBNext® UltraTM II Q5® Master Mix (NEB) as well as a range of Illumina adaptor primers (Merck) flanking the sgRNA cassettes. Following the purification of the PCR products using the PCR column purification kit (Qiagen) as well as the QIAquick gel extraction kit (Qiagen), they were multiplexed using q-PCR NEBNext library quant kit (E7630). 10nM of the multiplexed samples were then sequenced by next-generation sequencing using single-ended reads and data analysis was performed using DrugZ^72^.

### Clonogenic survival assays

Cells were seeded into 6-well plates at a concentration of 500-1500 cells/well in technical triplicate, with seeding density optimised to the cell line in question in order to account for baseline cell fitness defects where relevant (such as *TOPORS* KO cells with siRNF4). 24h after seeding, cells were treated with the indicated drugs or ionising radiation (IR) at the specified doses; formaldehyde-treated cells were subjected to drug washout and supplied with fresh growth medium after 24h of treatment. IR doses were performed using X-rays generated by a Rad Source RS 1800 Biological Irradiator. Six days after treatment, the surviving cells were fixed and stained with crystal violet. The number of colonies per well was counted, averaged between technical replicates and normalised to the number of colonies in untreated conditions. At least three biological replicates were performed per experiment, with each replicate displayed in quantification representing the mean normalised survival across three technical replicates.

For survival assays in Fig 5 e–g, cells were transfected with the corresponding siRNAs as described below. The transfected cells were reseeded into 6 well plates the following day (5,000 of HeLa cells or 10,000 of U2OS T-Rex cells per well). Cell confluency was monitored and analysed using IncuCyte S3 live cell imaging system every 12 hours for 5 days. After imaging, the cells were stained with crystal violet.

### Identification of proteins on nascent DNA (iPOND)

5×10^6^ HeLa TREx cells were seeded in two 15 cm dishes per condition and synchronised by double thymidine block. In brief, cells were seeded in the morning and thymidine containing media (2mM, T9250) was added after 8h. The next morning, cells were washed twice with 1x PBS and fresh, thymidine-free media was added for 9h before re-adding thymidine containing media and incubation overnight. Then, cells were washed twice with 1xPBS, released into thymidine-free media and treated with EdU (10μM) (Jena Bioscience, CLK-N001-100), 5-azadC (10μM) (Sigma, A3656), Ub-E1 inhibitor TAK-243 (1 μM) (Chemietek, AOB87172) or SUMO-E1 inhibitor ML-792 (5μM) (Axon Medchem, 3109) as indicated in figures. Treated cells were fixed in 10ml 1% FA for 20 min at room temperature followed. Unreacted FA was quenched by the addition of 1ml of 1.25M glycine and incubation at RT for 10 min. Cells were scraped into 50ml conical tubes, washed twice with 1x PBS, snap frozen in liquid nitrogen and stored at −80°C until processed. Technical duplicates were pooled at this step.

iPOND was essentially performed as described before^73^. Briefly, Pellets were resuspended in 1ml permeabilization buffer (0.25% Triton-X100 in 1xPBS) and incubated for 30 min at RT. The permeabilized cells were washed once with each 0.5% BSA in PBS and 1xPBS prior to resuspending in 500μl click reaction buffer (1xPBS, 10μM biotin-azide, 100mM sodium ascorbate, 100mM copper sulphate). After 1h incubation at room temperature, cells were washed once with each 0.5% BSA in PBS and 1xPBS and resuspended in 1 ml RIPA lysis buffer (100mM Tris pH 7.5, 150mM NaCl, 1% IGEPAL, 0.1% SDS, 0.5% sodium deoxycholate, cOmplete EDTA-free protease inhibitor cocktail). The lysate was sonicated in an ultrasonicator (Covaris E220 *evolution*) for 10 minutes followed by centrifugation at 21,130g for 10 min. The supernatant was incubated with 30μl streptavidin-sepharose (Cytavia, 90100484) at 4°C overnight. The next day, beads were washed with lysis buffer, 1M NaCl and once more with lysis buffer before snap freezing in liquid nitrogen and storage at −80°C.

### Identification of DNMT1-DPC proximal proteins by quantitative mass spectrometry

For quantitative mass spectrometry, beads were incubated for 1 hour in 2M Urea, 50mM Tris-HCl (pH 7.5), 5mM dithiothreitol containing trypsin. Beads were washed and the supernatant was saved. A reduction step with 5mM dithiothreitol and an alkylation step with 20mM chloroacetamide followed. Next, samples were digested overnight with trypsin. Samples were acidified using 1% formic acid and subsequently subjected to TMT labelling at 1.5:1 ratio for 1 hour in 150mM HEPES buffer (pH 8.5). TMT labelling was terminated with the addition of a 0.4% hydroxylamine solution, and excess labels were removed using reverse-phase Sep-Pak C18 cartridges. The TMT-labelled samples were pooled and desalted as previously described^74^. Peptide fractions were analysed on a quadrupole Orbitrap mass spectrometer (Orbitrap Exploris 480, Thermo Scientific) equipped with a UHPLC system (EASY-nLC 1000, Thermo Scientific) as described^75,76^. Peptide samples were loaded onto C18 reversed-phase columns (15 cm length, 75 μm inner diameter, 1.9 μm bead size) and eluted with a linear gradient from 8 to 40% acetonitrile containing 0.1% formic acid in 2 h. The mass spectrometer was operated in data-dependent mode, automatically switching between MS and MS2 acquisition. Survey full scan MS spectra (m/z 300–1650) were acquired in the Orbitrap. The 20 most intense ions were sequentially isolated and fragmented by higher-energy C-trap dissociation (HCD)^77^. Peptides with unassigned charge states, as well as with charge states less than +2 were excluded from fragmentation. Fragment spectra were acquired in the Orbitrap mass analyzer. Raw data files were analysed using MaxQuant (development version 1.6.14.0)^78^. Parent ion and MS2 spectra were searched against a database containing 98,566 human protein sequences obtained from UniProtKB (July 2021 release) using the Andromeda search engine^79^. Spectra were searched with a mass tolerance of 6 ppm in MS mode, 20 ppm in HCD MS2 mode and strict trypsin specificity, allowing up to three missed cleavages. Cysteine carbamidomethylation was searched as a fixed modification, whereas protein N-terminal acetylation and methionine oxidation were searched as variable modifications. The dataset was filtered based on posterior error probability (PEP) to arrive at a false discovery rate (FDR) of less than 1% estimated using a target-decoy approach^80^.

For statistical analysis, MaxQuant output data were imported into R. Only proteins identified in all four replicates of each condition were kept for downstream analysis. Intensities were log_2_ transformed and quantile normalised between the replicates using the R package preprocessCore. Significantly enriched proteins were identified by employing a moderated t-test using the R package limma with Benjamini-Hochberg FDR correction.

### Proximity Ligation Assay (PLA)

U2OS GFP-DNMT1 HA-EV or HA-TOPORS cells were seeded into 96-well imaging plates (PerkinElmer) at a density of 8×10^4^ cells per well. The following day, growth media was changed to normal growth media containing 2mM thymidine (Sigma-Aldrich) to arrest cells at the G1/S boundary. After 20-24h, cells were washed 4x in warm PBS, then washed once in normal growth media and incubated for 5 min at 37°C. Cells were then released into S-phase in normal growth media at 37°C in the presence or absence of 2µM SUMO E1i (Medchem Express) for 30 min. Media was then exchanged for treatment media containing 10µM 2’-deoxycytidine (dC; Sigma-Aldrich) or 5-aza-dC (Sigma-Aldrich) either with or without 2µM SUMO E1i and incubated at 37°C for 1h. Non-chromatin-bound proteins were cleared by pre-extraction in ice cold 0.2% Triton X-100 on ice for 2 min, after which cells were washed once with PBS and fixed with 4% formaldehyde for 15 min. PLA was then performed using anti-HA and anti-GFP antibodies; details of the antibodies used in this study can be found in Supplementary Table 3. The Duolink In Situ PLA Anti-Mouse Minus and Anti-Rabbit Plus probes (Sigma-Aldrich) and Duolink In Situ Detection Reagents FarRed Kit (Sigma-Aldrich) were then used as per the manufacturer’s instructions. Following PLA, cells were counterstained with 1µg/ml DAPI at room temperature for 2 min and then washed 4x with PBS. Images were acquired on a Zeiss 880 confocal microscope and analysis of PLA signal was performed using CellProfiler. Per-nucleus mean PLA intensity was calculated and normalised to the median per-nucleus mean PLA intensity of the GFP-DNMT1/HA-TOPORS PLA in 5-aza-dC-treated cells.

### Immunoprecipitations

For immunoprecipitation of GFP-TOPORS, five 15 cm dishes of HeLa cells per condition were transfected with the 5µg per dish of appropriate plasmids (plasmid details can be found in Table 2) and TransIT-LT1 (Mirus) according to the manufacturer’s instructions. The next day, cells were synchronized using 2mM Thymidine (Sigma-Aldrich). 20-24h later, cells were washed 4x with warm PBS and once in warm growth media and incubated for 5 min at 37°C. Cells were then released into normal growth media for 30 min at 37°C, then treated for 1h with 10µM dC or 5-aza-dC in the presence or absence of 2µM SUMO E1i, as indicated. Cells were washed twice with PBS, harvested by scraping in PBS and centrifuged at 300 x *g* for 3 min. Excess PBS was aspirated and cell pellets were snap-frozen on dry ice and stored at −80°C ahead of further processing.

Cells were lysed on ice for 10 min in ice cold IP150 buffer (50mM Tris-HCl pH 8, 150mM NaCl, 2mM MgCl_2_, 1% Triton X-100) supplemented with 500 U/ml benzonase (Sigma-Aldrich) and cOmplete EDTA-free protease inhibitor tablets (Roche). Cells were then sonicated using a chilled water bath sonicator for a total of 6 minutes with 30s ON/OFF pulses and placed on a rotor wheel at 4°C for 45 min. Lysates were then cleared by centrifugation in a bench-top microcentrifuge at full speed for 15 min at 4°C and standardised using IP150 buffer according to protein quantification by Bradford assay and 5% of each lysate was taken aside as an input sample, prepared for Western blot by boiling in 1X Laemmli buffer with 5% beta-mercaptoethanol and stored at −20°C. GFP-trap agarose beads (Chromotek) were equilibrated by washing twice in IP150 buffer with centrifugation at 0.3 x *g* for 2 min at 4°C. Equal amounts of equilibrated beads were added to each tube and samples were placed on a rotor wheel at 4°C overnight. Samples were washed 4 times by centrifuging for 1 min at 0.3 x *g* at 4°C, aspirating the supernatant and resuspending with 1ml ice cold IP150 buffer. Immunoprecipitates were eluted by boiling at 95°C for 5 min in 2X Laemmli buffer supplemented with 5% beta-mercaptoethanol.

### siRNA transfections

For PxP experiments, cells were seeded, 3×10^6^ cells in 10 cm dishes, in the morning and thymidine-containing media (2mM, T9250, Sigma) was added after 8h. After thymidine addition, 10µl siRNA (20µM) and 25µl Lipofectamine RNAiMAX Transfection Reagent (13778075, Thermo Scientific) were each diluted in 500µl Opti-MEM Medium. Following a 5 min incubation, siRNA and Lipofectamine RNAiMAX Transfection Reagent dilutions were mixed. After an additional 15 minutes, the transfection mix was added to cells. The next morning, the thymidine media was washed twice with PBS, trypsinized, counted and split in 6 cm dishes. Thymidine media was added again in the evening after cell attachment. The next morning cells were washed twice with PBS and released into fresh media for 2h before adding fresh media containing 5-aza-dC (10µM). After 30 min incubation, 5-aza-dC containing media was removed, cells were washed with PBS and allowed to recover. Cells were scraped on ice at the indicated timepoints and the pellets were frozen at −80°C.

### Purification of x-linked proteins (PxP)

PxP to isolate DNMT1-DPCs was essentially performed as described before^38^. In brief, cells were harvested by scraping into ice cold PBS and either snap-frozen or directly used for PxP. The cell pellet was resuspended in PBS at a concentration of 2×10^4^cells/μl. 10μl of the cell suspension was directly lysed in 1x LDS (NP0007, Thermo Scientific) as input samples. The remaining suspension was mixed with an equal volume of low melt agarose (2% in PBS, 1613111, Bio-Rad) and directly cast into plug moulds. After solidification at 4 °C for 5 min the plugs were transferred into 1ml of cold lysis buffer (1x PBS, 0.5mM EDTA, 2% sarkosyl, cOmplete EDTA-free protease inhibitor cocktail (4693132001, Merck), 0.04mg/ml Pefabloc SC (11585916001, Merck) and incubated on a rotating wheel for 4h at 4°C. For electroelution, plugs were fit into the wells of 10-well SDS-PAGE gels (12%, 1.5mm Novex WedgeWell). Electroelution was carried out at 20mA per gel for 60 min. After electroelution, plugs were transferred to 1.5ml tubes and boiled with 40μl 1xLDS sample buffer and 10μl reducing agent at 95°C for 20 min.

### Western blotting

Cell pellets were lysed in 10mM Tris pH 7.5 and 2% SDS, quantified through Bradford assays (LifeTechnologies), according to the manufacturer’s protocol, and stored at −20°C. After boiling the standardised protein lysates in 1X Laemmli buffer with 5% beta-mercaptoethanol, 20μg of the lysates were loaded on NuPage 4-12% gradient Bis-Tris gels (Invitrogen) and run at 150V for 90 min. The resolved proteins were then transferred onto a nitrocellulose membrane, which was blocked for 1 hour in 5% milk in TBS-T (0.1% Tween-20 in TBS) and analysed by standard immunoblotting using the antibodies listed in Supplementary Table 3. All Western blot images shown were obtained using the LI-COR platform (Biosciences) or by chemiluminescence using the ChemiDoc MP imager (BioRad).

### RNA Extraction and RT-PCR

Total RNA was extracted from harvested cells using the RNeasy Mini Kit with on column DNase digestion (Qiagen), according to the manufacturer’s instructions. During the extraction protocol, RNase-free plastic ware and solutions were used. After RNA extraction, isolated RNA was quantified using a Nanodrop spectrograph, and stored at −80°C. cDNA was reverse transcribed from 1μg of RNA using SuperScript IV VILO Master Mix (Invitrogen), according to the manufacturer’s instructions. cDNA was stored at −20 °C. qPCR was performed using 1μl of cDNA, 10μL of 2× Fast SYBR Green Master mix (Thermo Fisher Scientific) and 500nM forward and reverse primers, in a final volume of 20μL. Primers were designed and ordered to span an exon-exon junction of the target genes (Sigma-Aldrich) and are detailed in Supplementary Table 2. qPCR analysis was performed on a QuantStudio 5 Real-Time PCR System (Thermo Fisher Scientific), in technical triplicates. Gene expression changes were calculated using the 2–ΔΔCt method^81^.

## Acknowledgements

Research in the lab of J.S. is funded by European Research Council (ERC Starting Grant 801750 DNAProteinCrosslinks), Alfried Krupp Prize for Young University Teachers awarded by Alfriend-Krupp von Bohlen und Halbach Stiftung, European Molecular Biology Organization (YIP4644), The Vallee Foundation, and Deutsche Forschungsgemeinschaft (DFG, German Research Foundation) - Project ID 213249687 - SFB 1064 and Project-ID 393547839 - SFB 1361.

Research in the Beli lab is funded by the Deutsche Forschungsgemeinschaft (German Research Foundation, DFG): Project-ID BE 5342/3-1, Project-ID 393547839 – SFB 1361, Project-ID 464588647 – SFB 1551.

The S.P.J. laboratory is supported by Cancer Research UK (CRUK) Discovery Award DRCPGM\100005, CRUK core grant A:29580 and ERC Synergy Award 855741 (DDREAMM). The lab was also supported by CRUK Programme grant C6/A18796 and core funding grants C6946/A24843 and WT203144 to the Gurdon Institute. C.J.C., M.S-C., N.G., A.W., G.Z.V. and S.R. were funded by CRUK Programme grant C6/A18796 and CRUK Discovery Award DRCPGM\100005; C.P.C. is funded by a CRUK studentship DRCPGM\100005; V.G., A.S-B. and G.D’A. by ERC Synergy Award 855741. S.P.J. receives a salary from the University of Cambridge. We thank Nicola Lawrence and Kay Harnish of the Gurdon Institute Imaging and Scientific Facilities, at the Gurdon Institute, University of Cambridge.

For the purpose of open access, we have applied a Creative Commons Attribution (CC BY) public copyright licence to any Author Accepted Manuscript version arising from this submission.

## Author contributions

C.J.C, M.J.G., C.S.P-C, P.W., A.W., A.S-B., H-Y.L., C.J.B., M.S-C., G.Z-V, G.D’A, S.L.R. and N.G. generated cell lines and performed experiments. M.J.G. and V.G performed bioinformatic analyses. P.B. supervised work by C.J.B.. C.J.C, M.J.G, C.S.P-C., J.S. and S.P.J. wrote the manuscript. C.J.C, J.S and S.P.J coordinated and supervised the project.

---

**Supplemental Figure 1.**
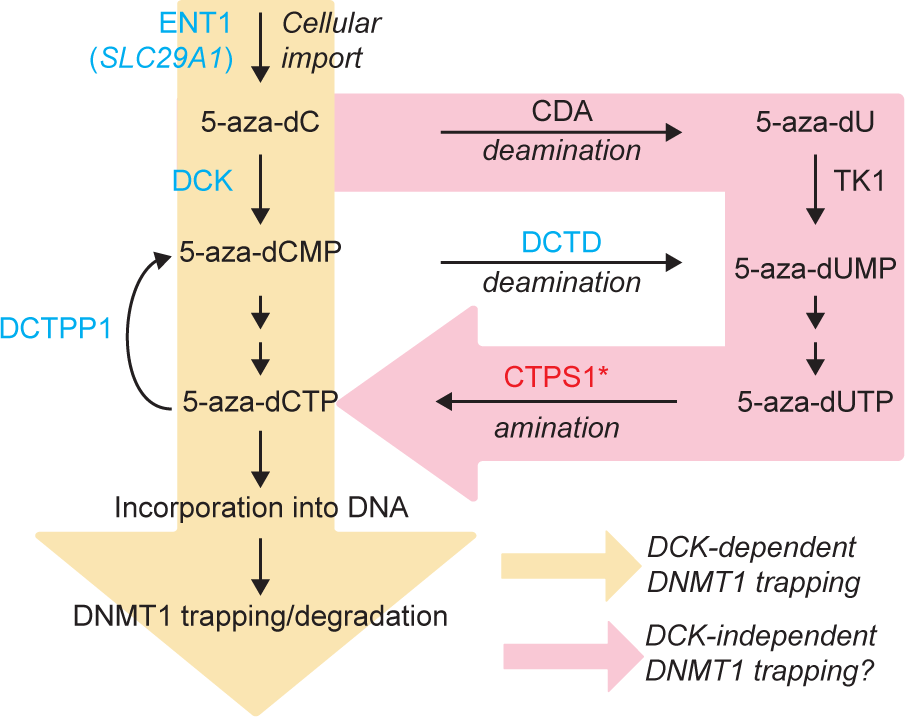
Speculative model for DCK-independent incorporation of 5-aza-dC into DNA and subsequent DNMT1 trapping. Briefly, upon cellular uptake, 5-aza-dC can be deaminated by CDA, followed by triphosphorylation involving the activity of TK1, followed by the possible conversion of 5-aza-dUTP to 5-aza-dCTP by CTPS1 and subsequent DNA incorporation and DNMT1 trapping.

**Supplemental Figure 2.**
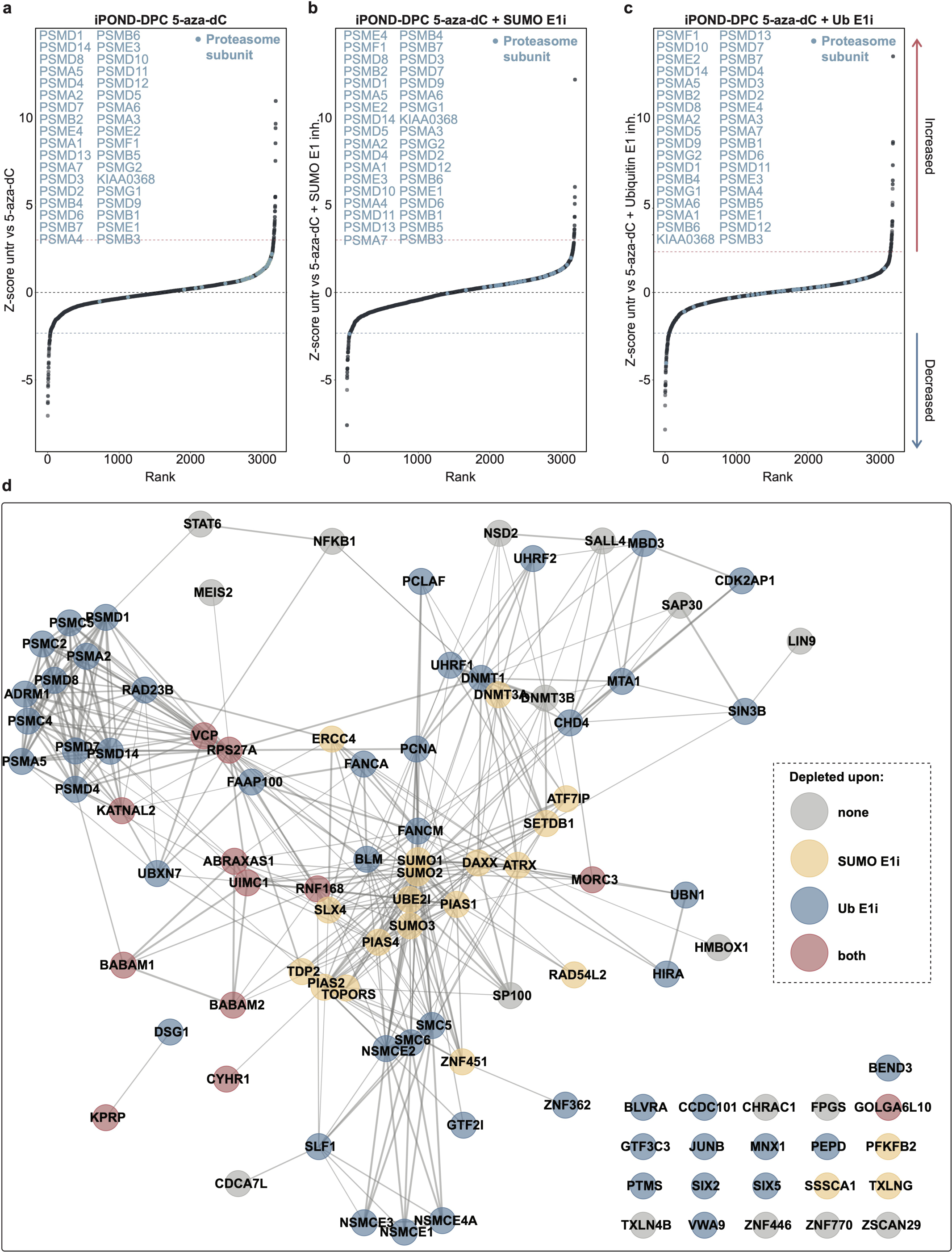
(a-c) Ranked standardised enrichment of proteasomal subunits detected by iPOND-DPC/MS from 5-aza-dC-treated (a), 5-aza-dC- and SUMO E1i-co-treated (b), and 5-aza-dC- and Ub E1i-co-treated (c) over untreated cells. (d) STRING analysis of proteins enriched on nascent DNA after 5-aza-dC treatment with SUMO/ubiquitin dependencies indicated, as assessed by iPOND-DPC/MS.

**Supplemental Figure 3.**
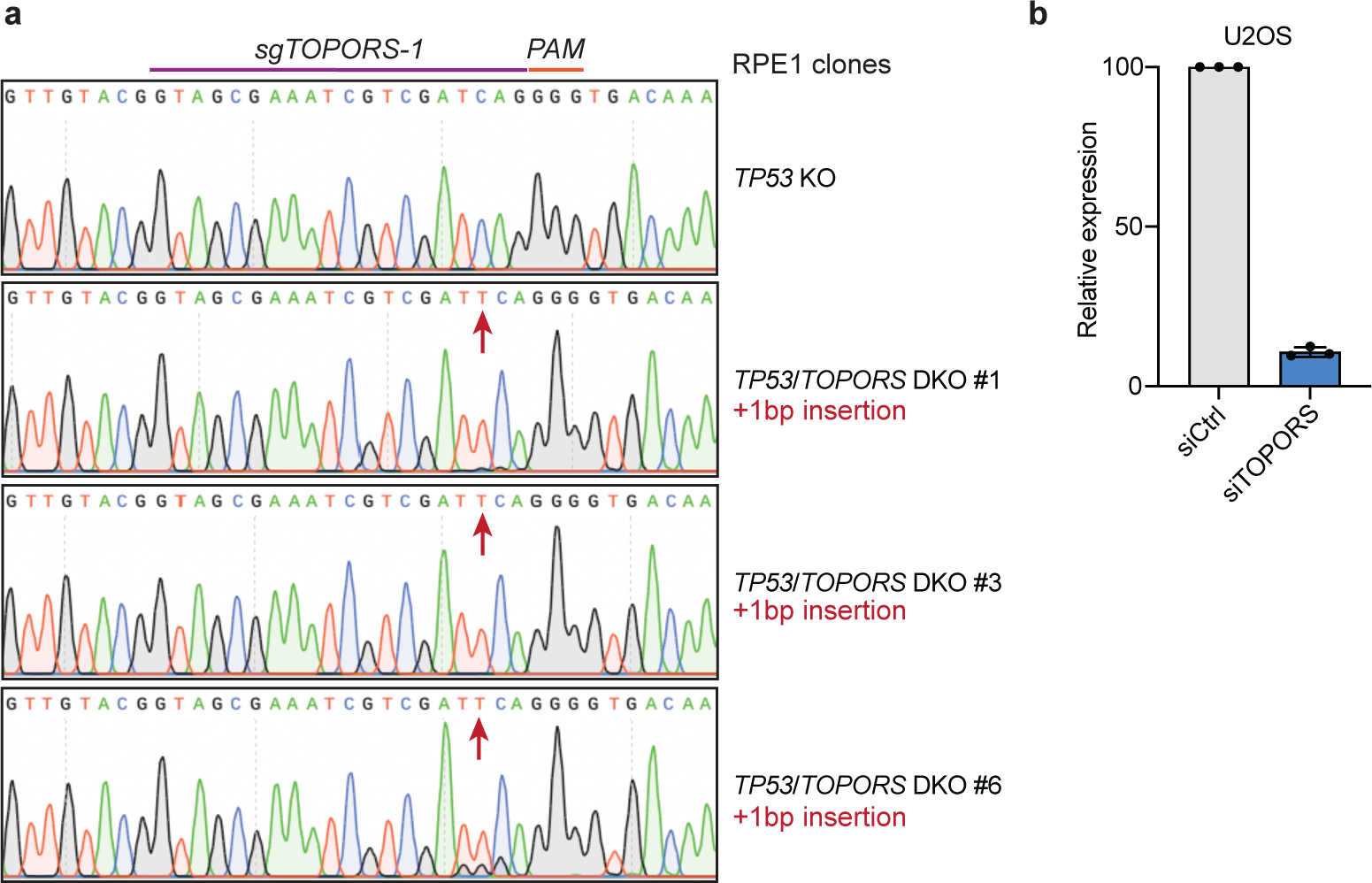
(a) Validation by Sanger sequencing of *TP53*/*TOPORS* DKO RPE1 clones. (b) Relative expression levels of TOPORS from U2OS cells 72h after siRNA-mediated depletion of TOPORS measured by qPCR, relative to GAPDH expression and normalised to TOPORS’ expression level siCtrl cells; n = 3 replicates, error bars ± SEM.

**Supplemental Figure 4.**
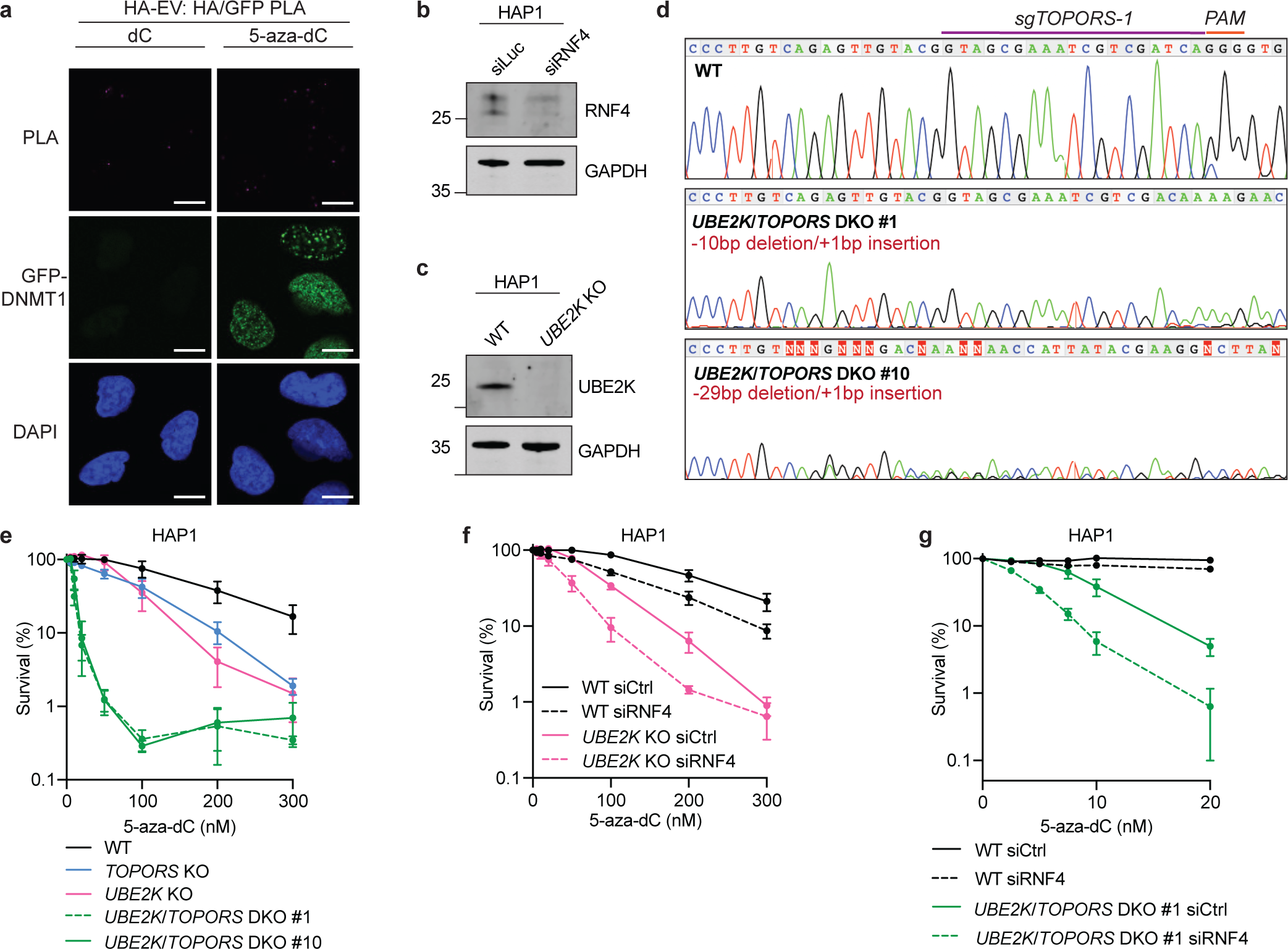
(a) Proximity ligation assay in U2OS cells expressing GFP-DNMT1 and HA-EV, treated with dC or 5-aza-dC. (b) Western blot of RNF4 from HAP1 cells after siRNA-mediated depletion of RNF4. (c) Western blot of UBE2K in WT and *UBE2K* KO HAP1 cells. (d) Validation by Sanger sequencing of *UBE2K*/*TOPORS* DKO HAP1 clones. (e) Clonogenic survival assays in WT, *UBE2K* KO, *TOPORS* KO and *UBE2K/TOPORS* DKO HAP1 cells treated with 5-aza-dC and stained after 6 days; n = 3 biological replicates, error bars ± SEM. (f) Clonogenic survival assays in WT and *UBE2K* KO HAP1 cells transfected with siCtrl or siRNF4, treated with 5-aza-dC 72h after transfection and stained 6 days later; n = 4 biological replicates, error bars ± SEM. (g) Clonogenic survival assays in WT and *UBE2K/TOPORS* DKO HAP1 cells transfected with siCtrl or siRNF4, treated with 5-aza-dC 72h after transfection and stained 6 days later; n = 3 biological replicates, error bars ± SEM.

